# Physiological and transcriptional changes in soybean as adaptive responses to the combined effects of soil alkalinity and drought

**DOI:** 10.1101/2024.08.30.610582

**Authors:** Md Rokibul Hasan, Asha Thapa, Mohammad Golam Mostofa, Ahmad H. Kabir

## Abstract

Soil alkalinity and drought collectively cause severe growth retardation in crops. However, the adaptive responses and transcriptional changes under such conditions remain unclear in soybean. In this growth incubator study, soil alkalinity and drought stress led to significant reductions in plant biomass, chlorophyll, and nutrient uptake in soybean. However, the photochemical efficiency of photosystem II remained stable, suggesting the activation of protective mechanisms to maintain photosynthetic functions. RNA-seq analysis demonstrated 357 upregulated and 799 downregulated genes in roots due to combined soil alkalinity and drought. Analysis revealed a complex response, with upregulation of genes predominantly involved in mineral homeostasis (*Iron dehydrogenase*, *Sulfurylase*), reactive oxygen species scavenging (*Glutamate synthase*, *L-Ascorbate peroxidase*) and hormonal signaling. Particularly, several ethylene-responsive genes, including the *Transcription factors TF5* and *TF018*, were upregulated, indicating the activation of stress-related signaling pathways. In a targeted study, plants supplemented with an ethylene precursor showed significant improvements in morpho-physiological traits and Fe status under combined stress. However, ethylene precursor applied in non-stressed plants led to reduced growth, and Fe levels, suggesting an involvement of the context-dependent role of ethylene in promoting stress tolerance. Furthermore, ethylene precursors caused an increase in root flavonoid and rhizosphere siderophore while restoring bacterial and fungal microbial cells in roots under combined stress in soil. However, in healthy plants, flavonoid and siderophore levels decreased, accompanied by a reduction in microbial cells to control levels. This suggests that elevated ethylene may regulate root exudates to recruit microbes dominated by host response, aiding soybean plants cope with combined stresses, although this effect may not occur in non-stressed plants. This is the first report on the transcriptional response and physiological adjustments in soybean exposed to alkalinity and drought, potentially advancing knowledge for genetic and agronomic interventions to improve stress resilience in legume crops.

## 1 Introduction

Global agricultural sustainability faces increasing challenges due to climate change, which exacerbates abiotic stressors such as drought and nutrient deficiencies. Soil alkalinity, characterized by high pH, is a critical factor that adversely affects plant growth and micronutrient availability (Xia et al., 2024). Elevated concentrations of sodium, carbonate, and bicarbonate in soils contribute to increased soil pH levels (Li et al., 2018). High pH conditions manifest in plants through various symptoms, including chlorosis, stunted growth, disease susceptibility, and altered coloration linked to impaired uptake of essential micronutrients such as iron (Fe), zinc (Zn), and manganese (Mn) (Samborska-Skutnik et al., 2020; Turner et al., 2020). Especially in legume crops, Fe is essential for nitrogen fixation and yield productivity (Briat et al., 2015). Consequently, alkalinity-induced micronutrient deficiencies significantly impair the structure and function of the photosynthetic machinery in plants (Singh et al., 2021a). Furthermore, drought stress reduces soil moisture and restricts mineral solubility, further negatively impacting crop yield (Sardans and Peñuelas, 2012; Waraich et al., 2011). Therefore, understanding how soil alkalinity and drought response in plants is pivotal for devising strategies to enhance crop resilience under multifaceted stress environments.

Plants employ various strategies to cope with soil alkalinity, enhancing their resilience and nutrient acquisition. One common approach is the release of organic acids from their roots, which help reduce the pH in the rhizosphere, thereby increasing the availability of essential nutrients (Barrow and Hartemink, 2023; Kabir et al. 2012). Morphologically, certain plant species develop extensive root systems to reach deeper soil layers and form mycorrhizal associations, which significantly enhance nutrient absorption (Zhang et al., 2020; Chen et al., 2016). Also, plant roots may release reducing or chelating substances (e.g., phenolics) into the rhizosphere, further mobilizing sparingly soluble micronutrients in alkaline soils (Wu et al., 2012). Furthermore, plants utilize various ion transporter genes, including H^+^-ATPases and Na^+^/H^+^ antiporters, to maintain ion homeostasis by regulating the uptake of essential cations and preventing toxic sodium accumulation (Kroh and Pilon, 2019; Assaha et al., 2017).

In plants, drought response involves intricate regulatory networks that coordinate responses at molecular, physiological, and morphological levels (Geng et al., 2024; Seleiman et al., 2021). In response to drought, plants possess a diverse array of adaptive strategies aimed at maintaining cellular hydration, sustaining metabolic processes, and minimizing damage during water deficit conditions. These strategies include morphological adjustments such as root architecture modification to enhance water uptake and leaf rolling to reduce water loss, alongside physiological mechanisms like osmotic adjustment, where compatible solutes accumulate to protect cellular structures (Volaire, 2018; Kumar et al., 2021; Khan et al., 2023; D’Oria et al., 2022). On a molecular level, plants undergo dynamic changes involving the activation of stress-responsive genes and signaling pathways. Among these, abscisic acid (ABA) signaling plays a central role by coordinating responses to water stress by stomatal closure to help the plant acclimate to drought conditions (Chaves et al., 2003; Ali et al., 2020). Additionally, ethylene contributes significantly to drought tolerance, not only by regulating drought-responsive genes but also by influencing cell wall remodeling and antioxidant responses, which are essential for minimizing oxidative damage under stress (Fatma et al., 2022). In addition, microbes also play a crucial role in stress resilience in plants. Flavonoids released by plants have been identified as pre-symbiotic signaling molecules in microbial-plant symbiosis (Lidoy et al., 2023; Schliemann et al., 2008). When plants are deficient in minerals, there is typically an increase in siderophores in the rhizosphere, secreted by both bacterial and fungal communities associated with host plants in a symbiotic relationship, to chelate Fe (Jin et al., 2010; Crowley et al., 2006).

Among staple crops, soybeans (*Glycine max*) play a crucial role in global food security and industrial applications (Kulkarni et al., 2018). Efforts to enhance soybean resilience to drought have yielded promising cultivars, yet their efficacy can be compromised under concurrent stress conditions, such as soil alkalinity. While the individual roles of soil alkalinity (Zocchi et al., 2017) and drought stress (Du et al., 2024) in limiting plant growth are well-documented, the interplay between soil alkalinity and drought tolerance remains underexplored. Particularly, the modulation of transcriptional changes and drought tolerance due to differential soil alkalinity in soybeans is still not well understood. Therefore, this study aims to unravel the physiological responses and transcriptome mechanisms governing the adaptive responses exclusively to soil alkalinity and drought tolerance in soybean. Furthermore, we examined the role of ethylene signaling and potential crosstalk with other hormones, providing insights into adaptive responses that may improve soybean resilience in response to combined soil alkalinity and drought in soybeans. These insights may optimize breeding strategies for developing resilient crop varieties of soybeans capable of withstanding complex environmental challenges.

## 2 Materials and methods

### 2.1 Plant cultivation and growth conditions

The seeds of soybean (var. Fiskeby IV) were collected from USDA-GRIN. Before germination, seeds were surface sterilized with 4% sodium hypochlorite solution for 5 min, followed by three washes with sterile water. Subsequently, seeds were placed in a tray and incubated at 25°C in the dark. Once uniform and healthy seedlings emerged in 3d, they were transplanted into soil pots (500 g soil), consisting of a mixture of natural field soil and commercial potting mix in a 1:2 ratio, for two distinct treatments: control (soil without lime, pH ∼6.5) and alkaline + drought (soil amended with 4.5 g NaHCO_3_ and 3.0 g CaCO_3_, pH ∼7.8). Throughout the cultivation period, 15 mM NaHCO_3_ was added weekly to maintain alkaline conditions (Kabir et al., 2012), while control plants received the same amount of water. Drought conditions were initiated by reducing water availability to 75% compared to the controls (well-watered) two weeks after transplanting the seedlings into the soil pots. Daily monitoring of soil moisture content was conducted to maintain 75% and 25% soil moisture levels for control and drought-stressed plants, respectively. Plants were cultivated following a randomized complete block design, with 9 replications (3 plants/pot) per treatment in a growth chamber (10 h light/14 h darkness, 250 μmol m ² s ¹) at 25°C up to 4 weeks before data measurement. In a targeted experiment, a precursor, 1-aminocyclopropane-1-carboxylic acid (ACC) of the ethylene synthesis (McDonnell et al., 2009), was supplemented in soil (10 mM) to validate whether it can induce tolerance in 5-week-old soybean plants in response to combined alkalinity and drought in soil.

### 2.2 Determination of morphological and photosynthetic parameters

The height of each plant’s shoot and the length of the longest root were measured using a digital caliper. The shoots and roots were then dried for three days at 75°C in an electric oven before their dry weights were recorded. Furthermore, chlorophyll scoring and chlorophyll fluorescence kinetics (OJIP) measurements, including Fv/Fm (maximal photochemical efficiency of PSII) and Pi_ABS (photosynthesis performance index) were conducted on the uppermost fully developed leaves at three different locations using a handheld chlorophyll meter (AMTAST, United States) and FluorPen FP 110 (Photon Systems Instruments, Czech Republic), respectively. For OJIP analysis, the leaves were acclimated to darkness for 1 h before data collection. Relative water content (RWC) of leaves was measured on the fully expanded trifoliate leaves at the tip of the main stem as previously described (Schonfeld et al., 1988). Briefly, the fresh weight (FW) of leaves from each treatment was recorded immediately after removal. The turgid weight (TW) was measured after soaking the leaves in distilled water in beakers for 24 h at room temperature. Following the soaking period, the leaves were quickly and carefully blotted dry with tissue paper to prepare for turgid weight determination. The dry weight (DW) of the leaves was obtained by oven-drying the leaf samples for 72 h at 75°C. RWC was calculated using the formula: RWC (%) = (FW – DW) / (TW – DW) x 100.

### 2.3 RNA-seq analysis in roots

RNA-seq analysis was performed on root samples collected from 4-week-old plants. The soybean roots were thoroughly cleaned by washing twice in sterile phosphate-buffered saline with 10 seconds of vortexing, followed by two rinses with sterile water. The cleaned roots were then ground into a powder with a pre-cooled pestle and mortar using liquid nitrogen. RNA was extracted from the powdered material using the SV Total RNA Isolation System (Promega, USA). For each sample, 1 μg of RNA with an RIN value above 8 was used for RNA-seq library preparations. The libraries were created with the KAPA HiFi HotStart Library Amplification Kit (Kapa Biosystems, USA) and sequenced on an Illumina NovaSeq 6000 instrument at the RTSF Genomics Core at Michigan State University. A total of 92.5% of reads passed filters with a quality score of ≥ Q30.

Bioinformatics analysis of raw FastQ files was performed using Partek Flow genomic analysis software (Partek, St. Louis, MO, USA). Adaptor sequences and low-quality reads were removed with Trimmomatic (Bolger et al., 2014) to obtain high-quality clean reads. The clean reads were aligned to the soybean reference genome, downloaded from HISAT2 (v2.0.5) (Yates et al., 2022; Kim et al., 2015). Gene read counts were determined using HTSeq (Anders et al., 2015). Differentially expressed genes (DEGs) were identified using the DESeq2 R package (Love et al. 2014), with FPKM (fragments per kilobase of exon per million mapped fragments) calculations. DEGs were defined as those with an adjusted P value < 0.05 and a fold change ≥ 2, using the false discovery rate (FDR) method. MA plot and heatmaps were created with ggplot2 and pheatmap (Hu et al., 2021) in R program, respectively. Annotation of DEGs was performed using Phytozome soybean database (Gmax_Wm82_a2_v1). Gene ontology and enrichment analyses of soybean DEGs were conducted using ShinyGO 0.81 (Ge et al., 2020).

### 2.4 Determination of tissue nutrient elements

Nutrient analysis was conducted on the roots and young leaves harvested from 4-week-old plants. The roots were detached, washed with running tap water, soaked in a 0.1 mM CaSO_4_ solution for 10 min, and then rinsed with deionized water. The young leaves were washed separately with deionized water. All samples were placed in envelopes and dried in an electric oven at 75°C for three days. Elemental concentrations were measured using inductively coupled plasma mass spectrometry (ICP-MS) at the Agricultural and Environmental Services Laboratories at the University of Georgia.

### 2.5 Estimation of flavonoid content in roots

The total flavonoid content in soybean roots was measured as described by Piyanete et al., (2009). In brief, 0.5 mL of 70% ethanol extract from the roots was mixed with 2 mL of distilled water and 0.15 mL of a 5% NaNO_2_ solution. After incubating for 6 min, 0.15 mL of a 10% AlCl_3_ solution was added and left to stand for another 6 min. Then, 2 mL of a 4% NaOH solution was added, and the mixture was diluted to a final volume of 5 mL with methanol and thoroughly mixed. After a 15-min incubation, the absorbance was measured at 510 nm using a spectrophotometer against a blank. The total flavonoid concentration was expressed in milligrams per gram of dry weight, calculated using a quercetin standard curve.

### 2.6 Determination of siderophore in the rhizosphere

We quantified the siderophore content in the rhizosphere soil using a chrome azurol S (CAS) assay, following the procedure outlined by Himpsl and Mobley (2019). Briefly, the rhizosphere soil was collected by washing off the soil tightly attached to the root surface. The rhizosphere soil was then homogenized with 80% methanol and subsequently centrifuged for 15 min at 10,000 rpm. The resulting supernatant (500 μl) was mixed with 500 μl of CAS solution and allowed to incubate at room temperature for 5 min. The absorbance of each reaction was then measured at 630 nm against a reference containing 1 ml of CAS reagent alone. The siderophore level per gram of soil sample was calculated using the formula: % siderophore unit = [(Ar - As) / Ar] x 100, where Ar represents the absorbance at 630 nm of the reference (CAS reagent only), and As represents the absorbance at 630 nm of the sample (CAS reagent + soil sample).

### 2.7 Determination of relative abundance of microbial cells

Harvested roots were cleaned by vortexing in sterile phosphate-buffered saline (PBS) for 10 sec, followed by two rinses with sterile water to remove surface contaminants. Approximately 0.2 g aliquots of cleaned root samples were used for DNA extraction following the procedure outlined by Clarke (2009), along with RNase and Proteinase K treatments. The DNA samples were then quantified and diluted before performing qPCR in a CFX96 Real-Time PCR Detection System (Bio-Rad, USA) using specific primers for 16S bacteria (16S-341_F: CCTACGGGNGGCWGCAG, 16S-341_R: GACTACHVGGGTATCTAATCC) and ITS fungi (ITS1-F: TCCGTAGGTGAACCTGCGG, ITS4-R: TCCTCCGCTTATTGATATGC). The soybean *Beta-tubulin* gene (Fw: GCTCCCAGCAGTACAGGACTCT, Rv: TGGCATCCCACATTTGTTGA) served as the internal control for relative quantification.

### 2.8 Data analysis

All data were collected from three independent biological replicates of individual plants for each treatment group. The statistical significance of each variable was assessed using Student’s *t*-test and one-way ANOVA, followed by Duncan’s Multiple Range Test (DMRT) where applicable. Graphical presentations were prepared using the ggplot2 package in R program.

## 3 Results

### 3.1 Changes in morphological traits and photosynthetic features

At weeks 3 and 4, we measured the morphological and photosynthetic attributes to study the responses of plants exposed to combined soil alkalinity and drought. The plants grown under soil alkalinity and drought showed a significant decrease in root length, root dry weight, shoot height, and shoot dry weight compared to the untreated controls (Figure 1A–1D). We observed a significant decrease in leaf SPAD score (an indicator of chlorophyll content) under alkaline conditions combined with drought compared to the controls (Figure 2A) at both time points. Fv/Fm remained the same in the alkaline and drought-stressed plants compared to the controls (Figure 2B). Interestingly, Pi_ABS showed no significant changes at week 3 but was significantly decreased under soil alkalinity combined with drought compared to the controls at week 4 (Figure 2C). We did not observe any significant changes in the relative water content (%) in the leaves of plants exposed to combined soil alkalinity and drought compared to the control at either 3 weeks or 4 weeks (Figure 2D).

**Figure 1.**
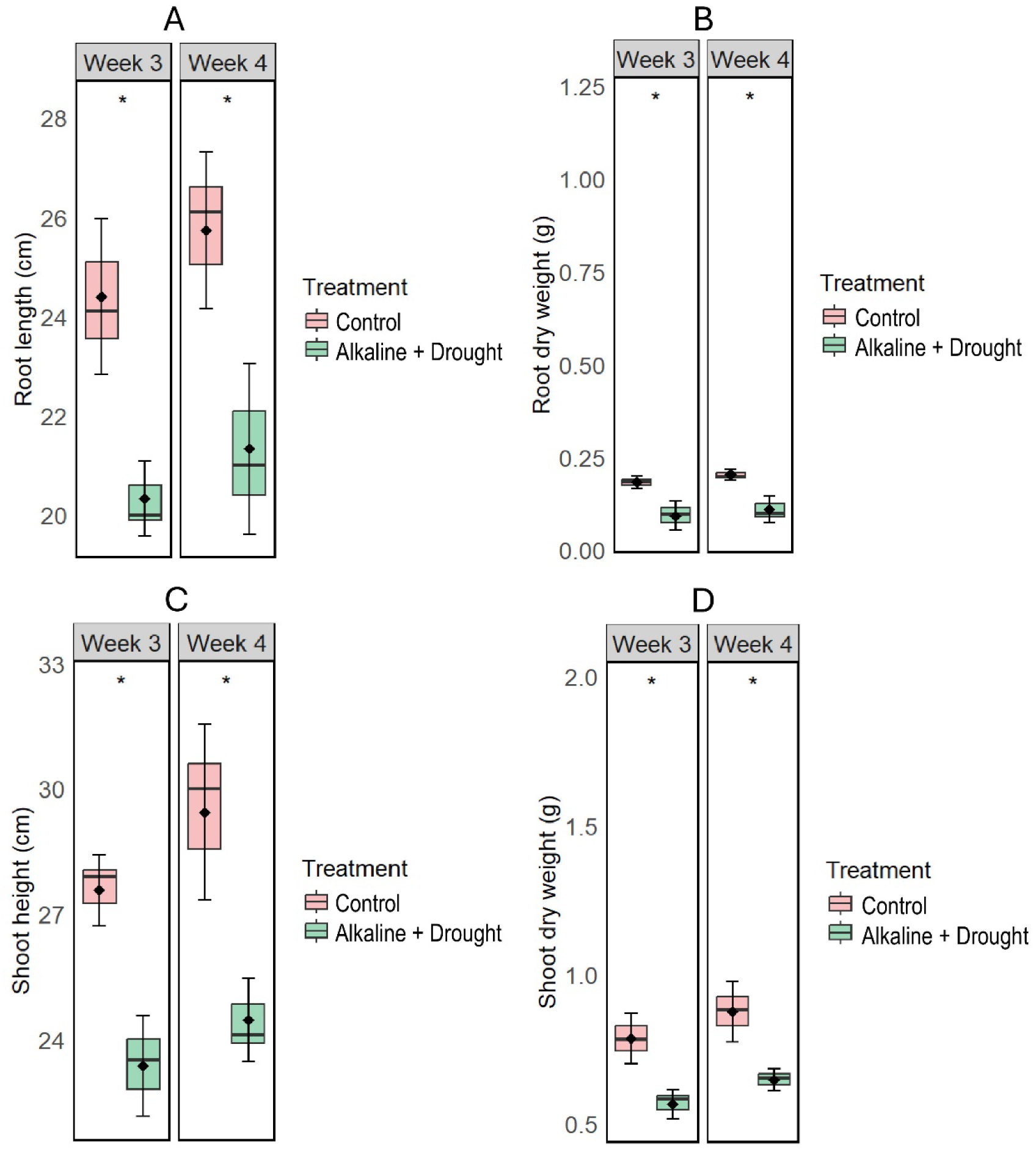
Changes in root length (A), root dry weight (B), shoot height (C), and shoot dry weight (D) in soybean cultivated for 3-week and 4-week in ‘control’ and ‘alkaline + drought’ conditions. Statistical significance between treatments was determined using a two-tailed *t*-test (* indicates significant differences, *p* < 0.05) The data represents the means ± SD of three independent biological samples (*n* = 3).

**Figure 2.**
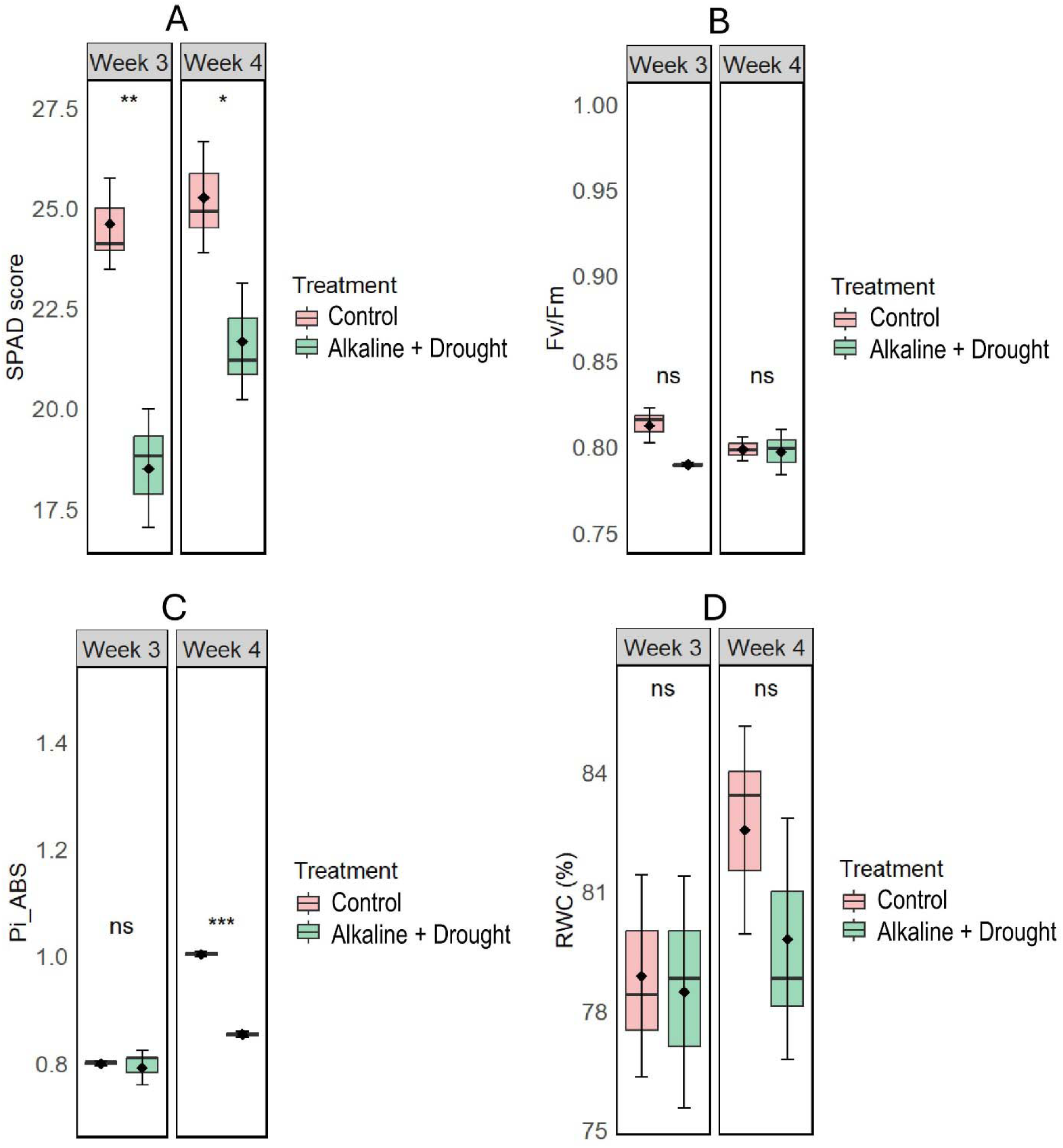
Changes in chlorophyll score (A), Fv/Fm: maximal quantum yield of PS II (B), and Pi_ABS: photosynthesis performance index (C) in the leaves of soybean cultivated for 3-week and 4-week in ‘control’ and ‘alkaline + drought’ conditions. Statistical significance between treatments was determined using a two-tailed *t*-test (\*\**p* < 0.01, \**p* < 0.05, \*\*\**p* < 0.001, and “ns” indicates no significant difference). The data represents the means ± SD of three independent biological samples (*n* = 3).

### 3.2 Elemental changes in plant tissues

The ICP-MS analysis of soybean roots and shoots under control and stress conditions revealed significant differences in nutrient concentrations (Table 1). Fe and N levels significantly decreased in both root and shoot under combined alkaline and drought conditions compared to controls. Zn, Mn, and Ca levels showed no significant changes in the roots due to soil alkalinity and drought conditions, but these minerals significantly declined in the shoot in such conditions relative to controls (Table 1). S and K levels significantly declined in the roots of soybean cultivated in response to combined alkalinity and drought compared to controls, while these minerals showed no significant changes in the shoot in such conditions. However, Mg levels showed no significant changes either in the root or shoot under soil alkalinity and drought conditions relative to untreated controls (Table 1).

**Table 1.**
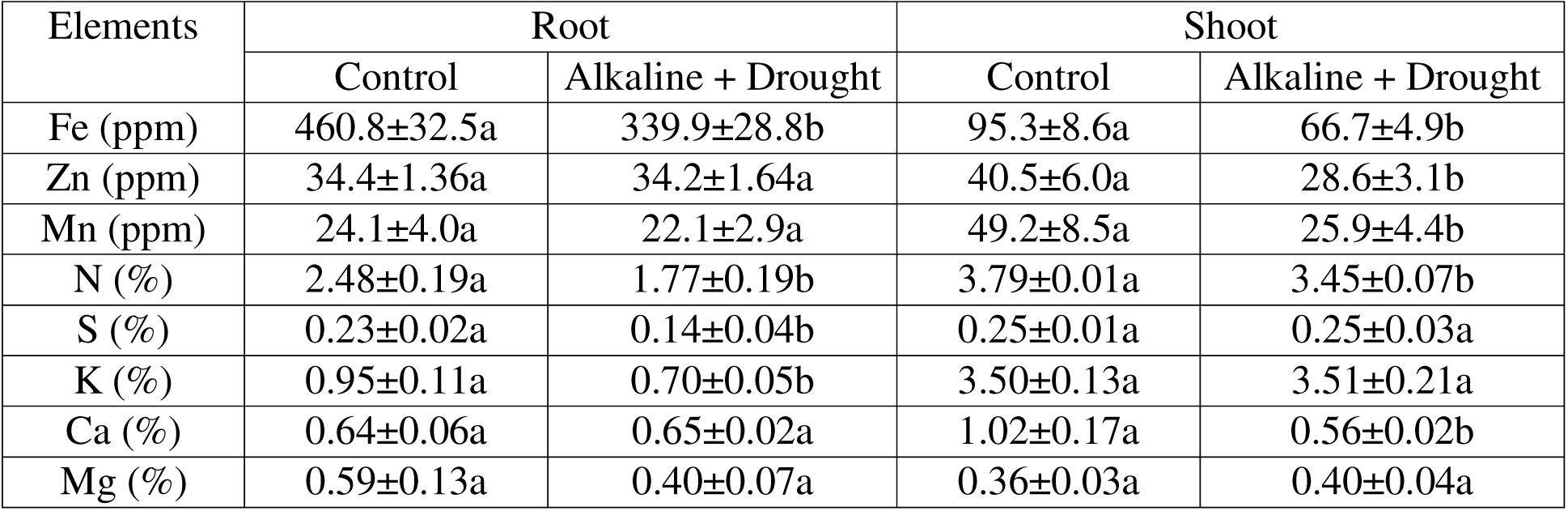
ICP-MS analysis of nutrients in the root and shoot of soybean grown in control and alkaline + drought conditions. Different letters indicate statistically significant differences between the treatment groups, as determined by a *t*-test at the *P* < 0.05 level. The data represents the means ± SD of three independent biological samples (*n* = 3).

| Elements | Root |  | Shoot |  |
| --- | --- | --- | --- | --- |
|  | Control | Alkaline + Drought | Control | Alkaline + Drought |
| Fe (ppm) | 460.8±32.5a | 339.9±28.8b | 95.3±8.6a | 66.7±4.9b |
| Zn (ppm) | 34.4±1.36a | 34.2±1.64a | 40.5±6.0a | 28.6±3.1b |
| Mn (ppm) | 24.1±4.0a | 22.1±2.9a | 49.2±8.5a | 25.9±4.4b |
| N (%) | 2.48±0.19a | 1.77±0.19b | 3.79±0.01a | 3.45±0.07b |
| S (%) | 0.23±0.02a | 0.14±0.04b | 0.25±0.01a | 0.25±0.03a |
| K (%) | 0.95±0.11a | 0.70±0.05b | 3.50±0.13a | 3.51±0.21a |
| Ca (%) | 0.64±0.06a | 0.65±0.02a | 1.02±0.17a | 0.56±0.02b |
| Mg (%) | 0.59±0.13a | 0.40±0.07a | 0.36±0.03a | 0.40±0.04a |

### 3.3 RNA-seq analysis of differentially expressed genes

We conducted RNA-seq analysis to examine the changes in the transcriptome of soybean roots harvested from 4-week-old plants exposed to combined soil alkalinity and drought. Principal component analysis (PCA) using three biological replicates revealed that the first two principal components, PC1 and PC2, accounted for 53.4% and 23.6% of the variance, respectively (Figure 3A). The separation between PC1 and PC2 indicates distinct grouping patterns among the samples, suggesting that treatment groups led to substantial differences in gene expression. Furthermore, we identified 357 upregulated and 799 downregulated genes differentially expressed under soil alkalinity and drought conditions compared to the control (Figure 3B).

**Figure 3.**
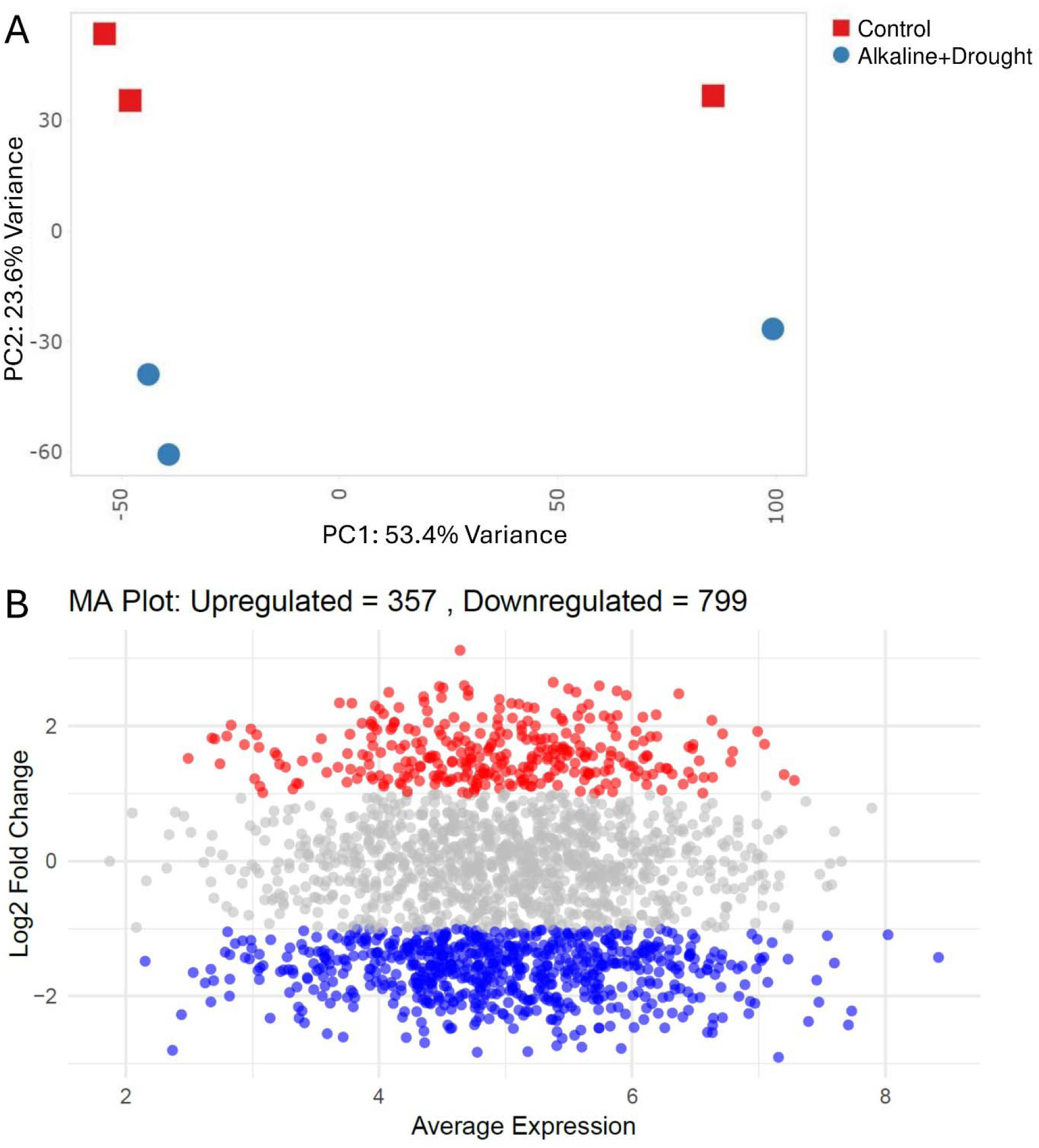
Analysis of RNA-sequencing data in roots of soybean cultivated for 4-week for differentially expressed genes (DEGs) in roots: A) principal component analysis of DEGs, B) MA plot of upregulated and downregulated genes (>Log2 fold, *P* < 0.05) in the roots of soybean grown in ‘alkaline + drought’ vs. ‘control’ conditions.

### 3.4 Heatmap analysis of candidate genes in roots

From the RNA-seq analysis, we present the top upregulated and downregulated candidate genes that are differentially expressed (> Log2 fold, < FDR value 0.05) and associated with different categories (Figure 4A, 4B). Under alkaline and drought stress, several genes were upregulated compared to the control related to mineral homeostasis which includes *Glyma.07G251300* (Fe dehydrogenase), *Glyma.20G151500* (Sulfurylase), and *Glyma.03G100000* (Copper ion transporter). Within hormonal signaling, *Glyma.07G215500* (ABA-glucosyltransferase), *Glyma.17G047300* (Ethylene-responsive TF18), *Glyma.13G088100* (Ethylene-responsive TF5), and *Glyma.19G181900* (Auxin response factor) were upregulated (Fig. 4A). In the stress response category, *Glyma.19G130800* (Glutamate synthase), *Glyma.12G073100* (L-Ascorbate peroxidase 2), *Glyma.02G261700* (Dehydration-responsive binding protein 2A), and *Glyma.10G227400* (ABC transporter C family 10) showed higher expression levels in the roots of soybean exposed to combined alkalinity and drought in soil (Fig. 4A). In addition, several genes related to mineral and solute transport such as *Glyma.10G146600* (Solute carrier family 40), *Glyma.17G203500* (Molybdate transporter 1), *Glyma.06G222400* (Amino acid transporter), *Glyma.02G305300* (Calcium-transporting ATPase), *Glyma.02G160500* (Copper transport protein), *Glyma.13G162900* (Anion transporter 6), and *Glyma.07G006500* (Sulfate transporter 3.4) were downregulated due to combined stress relative to controls (Fig. 4B). Within redox homeostasis, *Glyma.10G240300* (Peroxidase 66), *Glyma.10G193500* (Superoxide dismutase 2), *Glyma.01G163100* (Peroxidase 64), and *Glyma.04G019500* (Multi-copper oxidase type 1) were downregulated. In the category of pathogen immunity, Glyma.11G140800 (Thaumatin) and Glyma.14G070500 (Thioredoxin H1) showed decreased expression levels in response to combined soil alkalinity and drought (Figure 4B).

**Figure 4.**
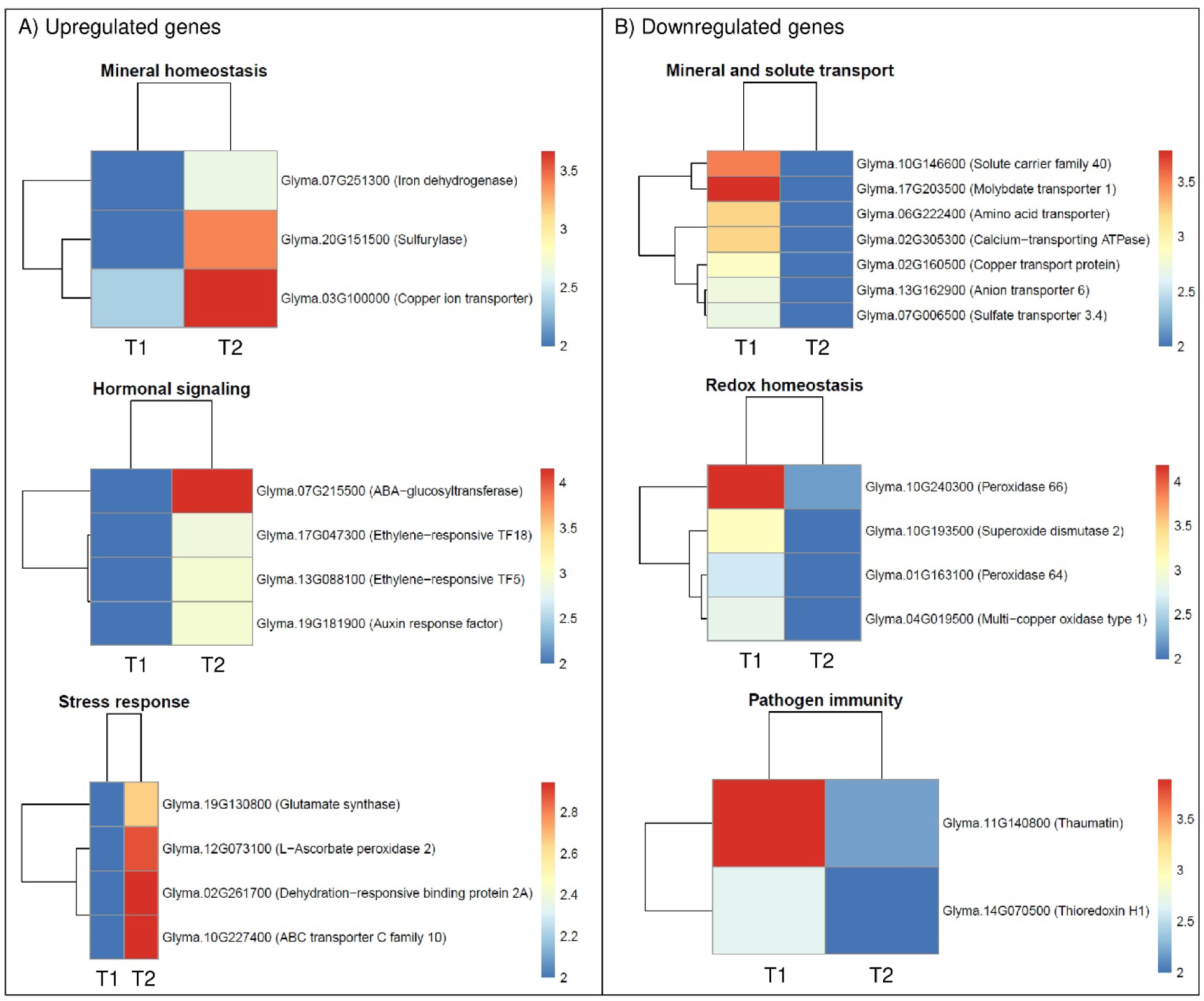
Heatmap of the selected differentially expressed genes (Log2 fold change > 1, *P* < 0.05) in the roots of soybean cultivated for 4-week in ‘alkaline + drought’ vs. ‘control’ conditions: A) Upregulated genes, B) Downregulated genes.

### 3.5 Gene enrichment analysis

Gene enrichment analysis revealed the enrichment of several pathways underlying the upregulated and downregulated genes (Figure 5A-5B). The upregulated genes showed significant enrichment in major molecular function, including sulfate adenylyltransferase (ATP) activity, glutamate synthase activity, oxidoreductase activities acting on CH–NH2 group donors, 5-methyltetrahydropteroyltriglutamate-homocysteine S-methyltransferase activity, ligase activity, phospholipid-translocating ATPase activity, peptidase activity, and peptidyl-prolyl cis-trans isomerase activity, alongside Fe ion and heme binding (Figure 5A). In the KEGG pathway analysis, the upregulated genes showed enrichment in pathways including phosphonate and phosphinate metabolism, selenocompound metabolism, monobactam biosynthesis, nitrogen metabolism, sulfur metabolism, cyanoamino acid metabolism, phenylalanine, tyrosine, and tryptophan biosynthesis, biosynthesis of plant secondary metabolites, terpenoid backbone biosynthesis, glyoxylate, and dicarboxylate metabolism, biosynthesis of amino acids, biosynthesis of secondary metabolites, and metabolic pathways (Figure 5A). Furthermore, the molecular function analysis of downregulated genes showed significant enrichment in ferric Fe binding, DNA-directed DNA polymerase activity, 3’-5’ exonuclease activity, calmodulin binding, Fe-sulfur cluster binding, phosphotransferase activity with alcohol group as acceptor, nucleotide binding, electron carrier activity, and metal ion binding (Figure 5B). In the KEGG pathway analysis, the downregulated genes were enriched in pathways including caffeine metabolism, thiamine metabolism, autophagy, DNA replication, phosphatidylinositol signaling system, inositol phosphate metabolism, nucleotide metabolism, phagosome, peroxisome, purine metabolism, and other metabolic pathways (Figure 5B).

**Figure 5.**
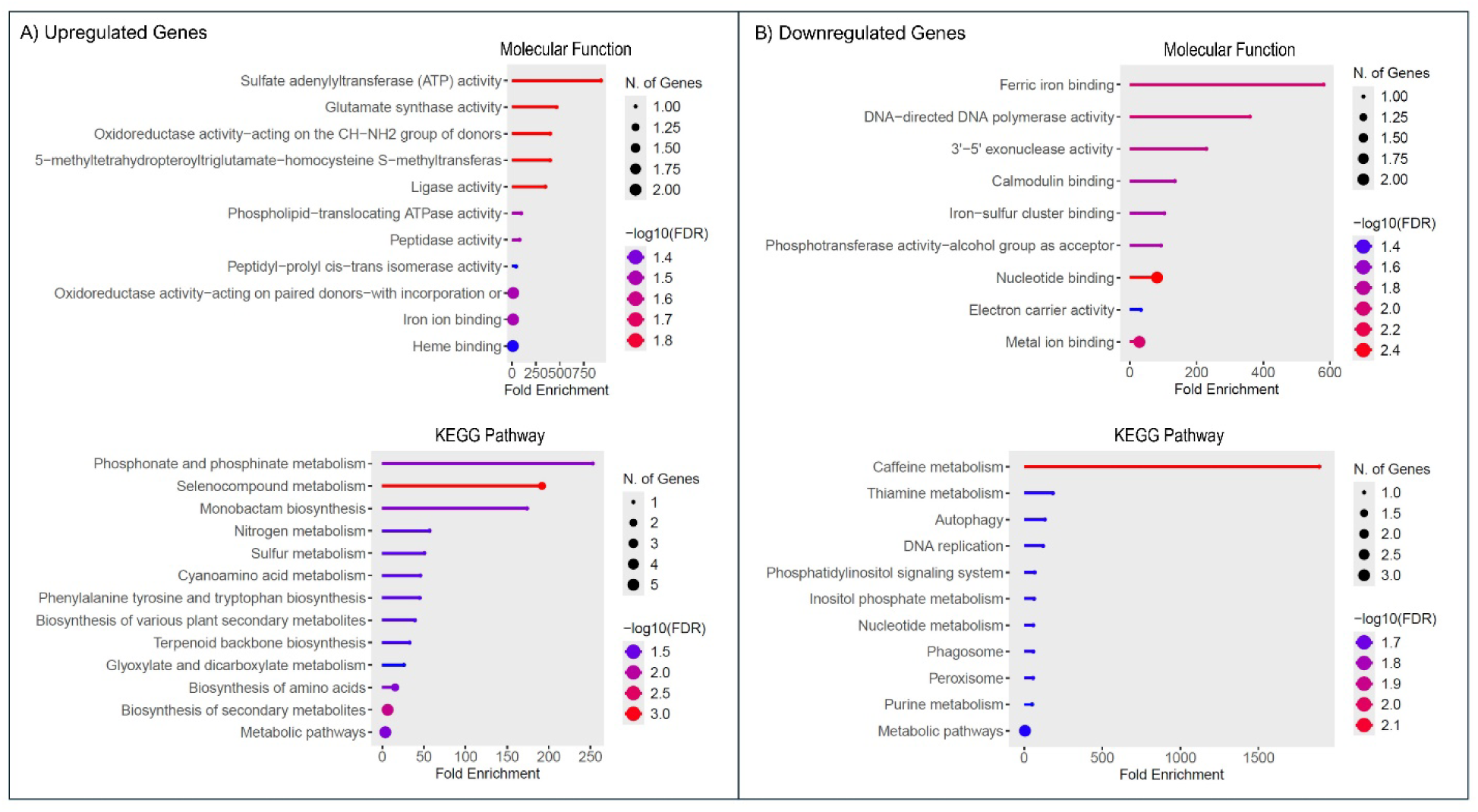
Gene enrichment analysis for the upregulated (A) and downregulated genes (B) in roots of soybean cultivated in ‘alkaline + drought’ and ‘control’ conditions for molecular function, and KEGG pathway. The size and color of the data points indicate the number of genes involved and the significance level, respectively, with larger points and deeper colors representing more substantial enrichment and higher significance.

### 3.6 ACC (ethylene precursor) on promoting dual stress tolerance

Given that we observed the upregulation of several ethylene-responsive genes in RNA-seq, we further investigated whether ACC (ethylene precursor) has significant biological effects on recovering dual stress in soybean. In this study, the combined alkalinity and drought exhibited a significant decrease in chlorophyll score, plant height, plant dry weight, stem diameter, and leaf RWC (%) in soybeans relative to untreated controls (Figure 6A-6F). ACC supplementation resulted in significant improvements in these traits compared to the alkaline + drought conditions without ACC (Figure 6A-6F). Surprisingly, the addition of ACC under control conditions resulted in the lowest SPAD score, plant height, and plant dry weight (Figure 6B-6D). However, the stem diameter and leaf RWC (%) followed a similar trend to that observed in dual-stressed plants and controls, respectively (Fig. 6E-6F). ICP-MS analysis showed a significant decrease in leaf Fe concentration due to alkaline + drought conditions compared to controls. However, the addition of ACC under dual alkalinity and drought in soil caused a significant increase in Fe levels in the leaf compared to the plants in such conditions without ACC (Figure 6G). Furthermore, the ACC treated on control plants showed the lowest leaf Fe level to those of other treatment groups (Figure 6G). We found that alkaline + drought conditions showed no significant changes in flavonoid content in the roots of soybean relative to controls. However, the addition of ACC under soil alkalinity and drought caused a significant increase in flavonoid content compared to those plants solely cultivated under combined alkalinity and drought without ACC. Control plants supplemented with ACC showed similar flavonoid levels in the roots to those of control (Figure 6H). Interestingly, siderophore levels in the rhizosphere of soybean exposed to alkaline + drought conditions showed a significant increase relative to controls (Figure 6I). Soybean plants exposed to combined alkalinity and drought supplemented with ACC showed a significant increase in rhizosphere siderophore compared to those plants exposed to alkaline + drought conditions without ACC supplementation. Lastly, control plants supplemented with ACC showed similar rhizosphere siderophores to those of controls (Figure 6H).

**Figure 6.**
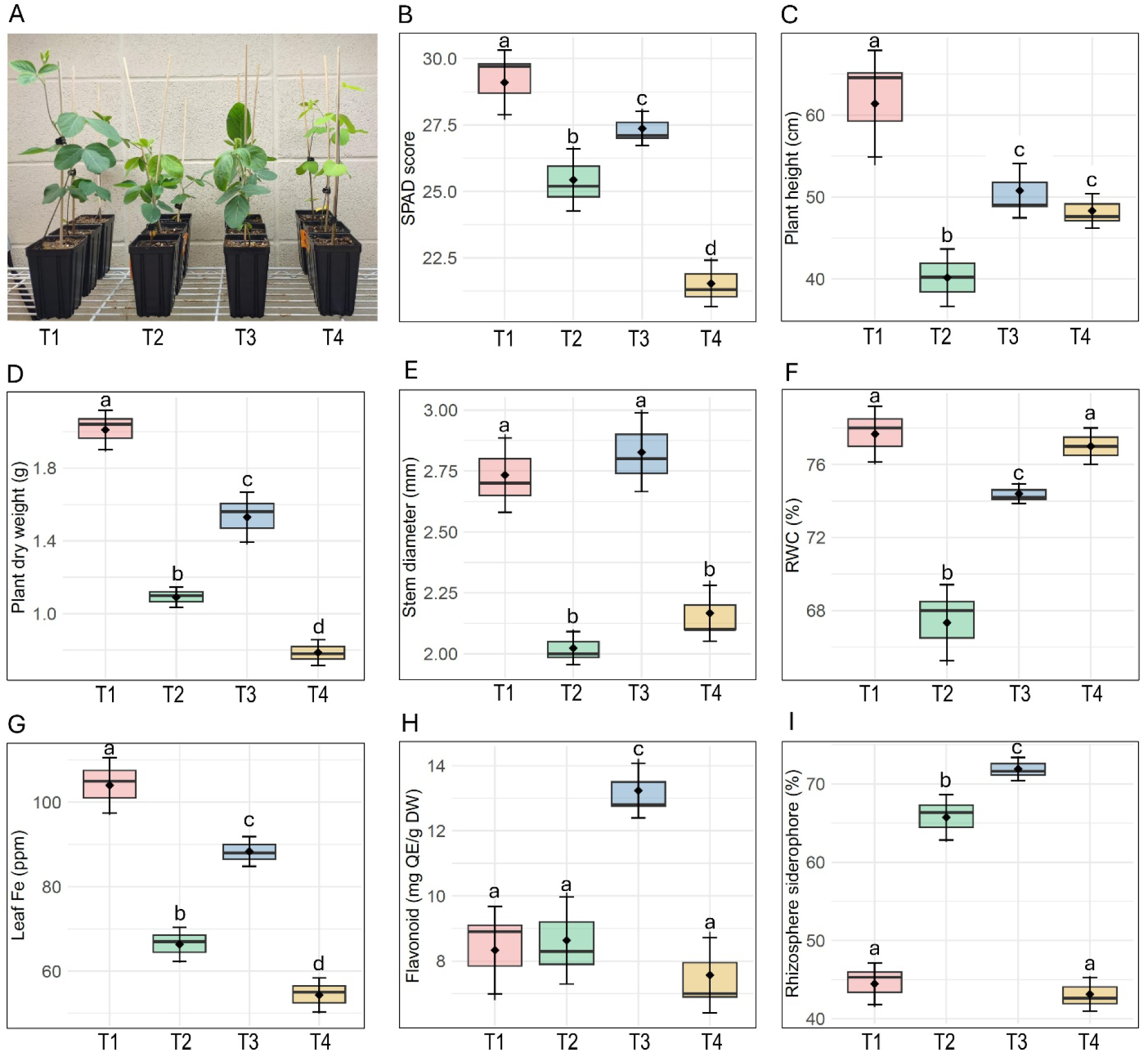
Changes in plant phenotype (A), chlorophyll score (B), plant height (C), plant dry weight (D), stem diameter (E), relative water content (F), leaf Fe (G), flavonoid content (H) and rhizosphere siderophore (I) in soybean exposed to different conditions (T1: control, T2: alkaline + drought, T3: alkaline + drought + ACC, T4: control + ACC) for 4 weeks. Different letters above the bar indicate significant differences between treatment groups at a *P* <0.05 level, where applicable. The data represents the means ± SD of three independent biological samples (*n* = 3).

We further determined the relative of 16S and ITS microbial cells in roots by DNA-based qPCR. We found that combined soil alkalinity and drought caused a significant decline in 16S bacterial abundance in roots relative to controls. However, the addition of ACC under soil alkalinity and drought conditions showed a significant increase in 16S, which is similar to controls. Control plants supplemented with ACC showed similar 16S levels in the roots to those of controls (Figure 7A). Similarly, the abundance of ITS fungal cells significantly decreased in the roots due to soil alkalinity and drought in contrast to untreated controls. Interestingly, ITS fungal abundance showed a significant increase when ACC was added to soil exposed to combined soil alkalinity and drought. Also, the increase in ITS abundance was relatively higher than the plants cultivated in control conditions. Furthermore, plants treated with ACC under control conditions showed similar ITS abundance in the roots to those of untreated controls (Figure 7B).

**Figure 7.**
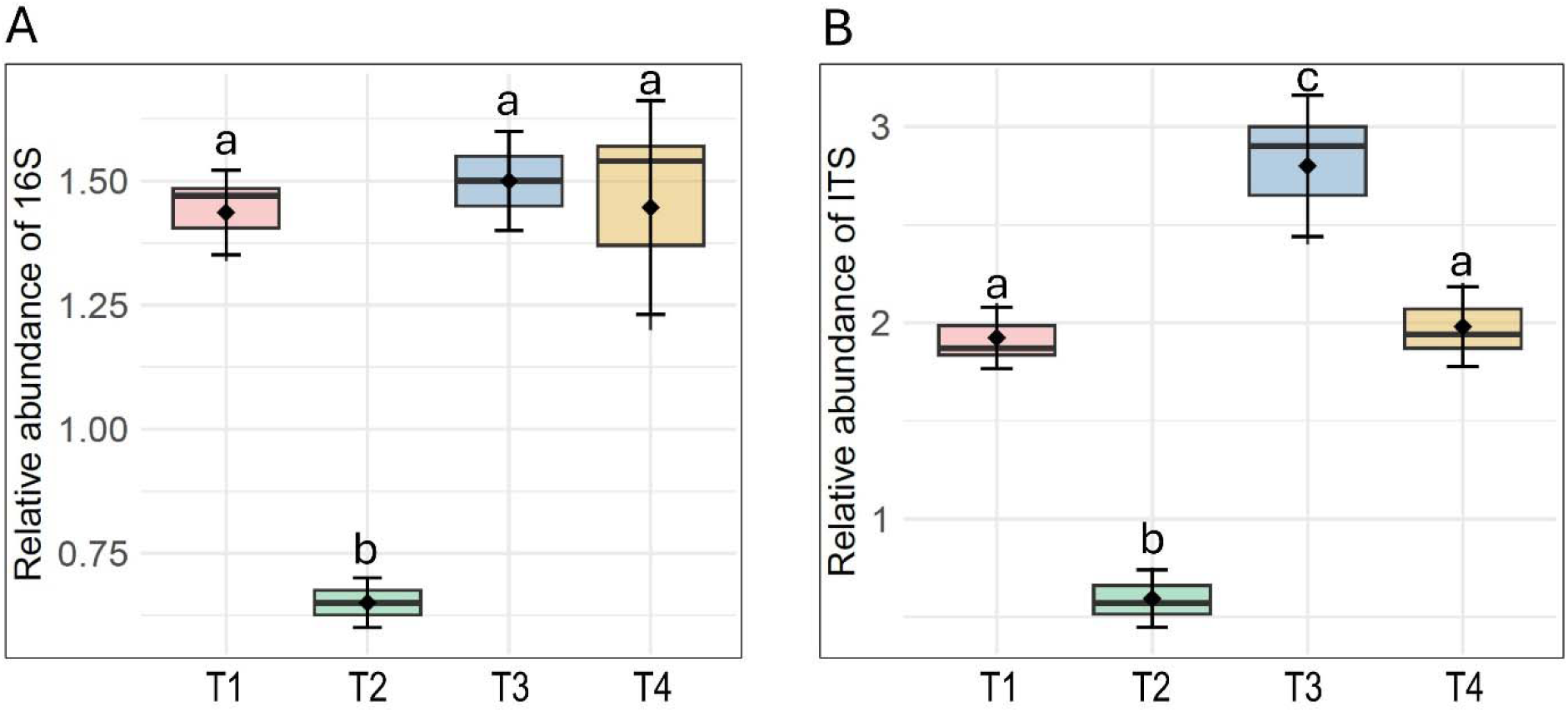
Changes in the relative abundance of 16S (bacteria) and ITS (fungi) cells in roots of soybean exposed to different conditions (T1: control, T2: alkaline + drought, T3: alkaline + drought + ACC, T4: control + ACC) for 4 weeks. Different letters above the bar indicate significant differences between treatment groups at a *P* <0.05 level, where applicable. The data represents the means ± SD of three independent biological samples (*n* = 3).

## 4 Discussion

This study reveals the physiological and transcriptional changes along with the induction of tolerance ability in soybean exposed to combined soil alkalinity and drought. Plants are constantly exposed to combined abiotic stresses, such as soil factors and drought, in their natural environments. These simultaneous challenges can lead to an intensified effect, making it harder for plants to adapt and survive (Teshome et al., 2020). This is the first growth incubator study to reveal transcriptional changes, with many genes related to growth and stress responses being differentially expressed in soybean stressed with alkalinity and drought in soil. We also observed the requirement for ethylene-mediated improvement in plant health and possible indications of symbiosis that may be necessary for combined stress tolerance in soybean.

The significant reduction in plant biomass subjected to alkalinity and drought stress, as observed in our study, indicates the detrimental impact of these combined abiotic stresses on the drought-tolerant soybean genotype. These morphological changes are likely due to impaired cellular processes and metabolic activities resulting from inadequate Fe availability and water stress. Fe is a critical micronutrient involved in various physiological processes, including chlorophyll synthesis and electron transport in photosynthesis (Samira et al., 2013). The observed decrease in chlorophyll content and photosynthetic performance index under alkaline and drought stress conditions underscores the compromised photosynthetic capacity of the plants. Interestingly, the maximum quantum efficiency of photosystem II (Fv/Fm) remained unaffected under stress suggesting that while the overall photosynthetic apparatus is impacted, the core photochemical efficiency of PSII remains intact. It is possible that the non-photochemical quenching helps dissipate excess light energy as heat to prevent damage to PSII reaction centers (Ruban et al., 2016; Demmig-Adams et al., 2014). Furthermore, enhanced scavenging of ROS may also contribute to the stability of PSII by neutralizing ROS generated under stress (Foyer & Noctor, 2011). We observed the upregulation of *L-Ascorbate Peroxidase 2* in the roots of soybean. *Ascorbate Peroxidase 2* reduces H O to water using ascorbate as an electron donor, playing a key role in stress tolerance and maintaining cellular redox homeostasis (Hong et al. 2018).

The elemental analysis of soybean tissues revealed significant alterations in nutrient concentrations due to alkaline and drought stress. The marked decrease in Fe and N levels in both roots and shoots highlights the impaired nutrient uptake and transport mechanisms under these stress conditions. Interestingly, we found the upregulation of *Glutamate synthase* and *Fe dehydrogenase* known for their role in nitrogen assimilation in the roots. It may be a compensatory response of soybean plants in nitrogen and Fe metabolism to combat stressed conditions. *Glutamate synthase* assimilates ammonia by catalyzing the synthesis of glutamate from glutamine and 2-oxoglutarate, a process vital for amino acid biosynthesis (Masclaux-Daubresse et al. 2006). However, the unchanged levels of Zn, Mn, and Ca in the roots, despite significant declines in the shoots under alkalinity and drought stress, suggest differential regulation of nutrient allocation and homeostasis between root and shoot tissues. The stability of these elements in the roots might be attributed to their relatively lower mobility and the plant’s efforts to prioritize root functionality under stress. Furthermore, the significant reduction in S and K levels in the roots, coupled with no significant changes in the shoots, further suggests the differential nutrient management strategies adopted by the plants under soil alkalinity and drought stress. S is vital for the synthesis of amino acids and proteins (Ren et al., 2022; Künstler et al., 2020). Furthermore, potassium regulates stomatal exposure, helping to reduce water loss through transpiration and maintain turgor pressure under drought conditions (Wang et al., 2013). K also activates numerous enzymes, including those involved in photosynthesis, protein synthesis, and carbohydrate metabolism, helping plants sustain metabolic functions under stress (Ho et al., 2022; Jin et al., 2011). Additionally, K is critical for maintaining ion homeostasis and enhancing the plant’s ability to uptake other essential nutrients (Lian et al., 2023). Hence, the retention of K levels in the shoot may therefore reflect the soybean plants’ attempt to optimize water usage and nutrient uptake under combined alkaline and drought stress. Furthermore, the retention of sulfur levels in the shoot aligns with the gene enrichment in oxidoreductase activity, indicating a significant role in redox reactions and sulfur metabolism in soybeans exposed to the combined effects of alkaline and drought in soil. Several studies demonstrated that redox reactions and sulfur metabolism play a critical role in responses to abiotic stress plants (Mittler, 2002; Kopriva and Rennenberg, 2004).

Furthermore, soybean plants exposed to concurrent soil alkalinity and drought showed upregulation of the *ABA-glucosyltransferase*. It is a major player in the ABA conjugation pathway for regulating ABA homeostasis and adapting to abiotic stresses (Dong and Hwang, 2014; Xu et al., 2002). Although ABA is a well-known stress hormone, particularly involved in drought responses, it is also known to induce alkaline stress tolerance in plants (Feng et al. 2024, Xu et al. 2022). This crosstalk between ABA and alkaline stress is particularly important in drought conditions, where the plant’s need for nutrient retention might increase due to compromised root functionality. Interestingly, we found the upregulation of several ethylene-responsive genes, such as ethylene-responsive transcription factors (TF5 and TF018) and dehydration-responsive binding protein 2A, in response to combined alkalinity and drought in soybean. Naing et al. (2022) showed that ethylene-insensitive mutants exhibited increased sensitivity to drought stress, indicating that ethylene-responsive genes contribute to drought adaptation by modulating stomatal conductance and enhancing root growth. This may imply the activation of stress-related signaling pathways and the role of ethylene signaling in soybean’s adaptive response to combined stress. Research indicates that ethylene regulates root growth inhibition caused by alkaline stress by promoting the expression of genes involved in auxin biosynthesis (Li et al. 2015). Similarly, we also found the induction of *Auxin response factor* (*Glyma.19G181900*) in the roots of soybean due to stress. Particularly, auxin plays a critical role in mediating adaptive responses in plants (Liu et al., 2024; Iqbal et al., 2022; Kabir et al., 2012). Additionally, ethylene biosynthesis may induce drought-responsive genes that help plants conserve water and maintain cellular homeostasis (Mohorović et al., 2023; Arraes et al., 2015). Ethylene signaling may also facilitate ROS scavenging enzymes, helping mitigate oxidative stress in plants (Zhang et al., 2016; Kabir et al., 2012). In most cases, ethylene, whose production increases under nutrient deficiency in genotypically efficient genotypes, acts as an activator of stress responses in plants (Dubois et al., 2018). Previous studies indicating the upregulation of *1-aminocyclopropane-1-carboxylic acid synthase (ACS)*, and *1-aminocyclopropane-1-carboxylic acid oxidade* genes involved in ethylene synthesis further support the increased ethylene production as part of the adaptive responses under lime-induced Fe deficiency in plants (García et al., 2015; Sauter et al., 2013). Thus, it appears that ethylene-related genes could be potential targets for improving stress tolerance in soybean exposed to concurrent soil alkalinity and drought.

To further elucidate the role of ethylene, we conducted a targeted study using an ethylene precursor (ACC) to determine if it could eliminate the impairment in soybean exposed to combined alkalinity and drought. In this study, supplementation with ACC led to significant improvements in these morpho-physiological traits, suggesting that ethylene signaling helps restore key growth parameters along with leaf Fe status disrupted by combined stress conditions. Surprisingly, the addition of ACC to control plants caused a drastic decrease in the growth parameters of soybean, especially in leaf chlorophyll and Fe levels. This response may be attributed to the disruption of hormone homeostasis in plants under non-stress conditions. Elevated ethylene levels in optimal conditions can induce premature senescence leading to chlorophyll degradation and reduced Fe uptake (Koyama, 2018; Kazan, 2015). Therefore, the contrasting effects of ACC under stressed versus non-stressed conditions in soybean underscore the context-specific role of ethylene signaling.

Interestingly, we observed a correlation between root flavonoid content and rhizosphere siderophore levels induced by the presence of an ACC, further underscoring the significance of microbial associations in soybean subjected to combined stress. While ‘alkaline + drought’ stress conditions alone did not affect root flavonoid content, ACC supplementation significantly increased flavonoid levels. This enhancement in flavonoids might contribute to improved stress resilience in soybean as flavonoids are known to play roles in symbiotic associations (Wang et al., 2022). Flavonoids play an essential role in rhizobium-legume symbiosis as chemoattractants and *Nod* gene inducers, and they are also believed to stimulate fungal spore germination and hyphal growth (Abdel-Lateif et al., 2012; Hassan et al., 2012). Similarly, siderophore levels in the rhizosphere were elevated under combined alkaline and drought conditions and ACC further boosted siderophore production. This indicates that ACC not only improves plant tolerance to concurrent stress but also potentially aids in Fe acquisition by increasing siderophore-mediated Fe mobilization in the rhizosphere (Figure 8). Microbial siderophores are compounds secreted by both bacterial and fungal microbial communities that bind and solubilize Fe, making them more accessible for uptake by plants (Singh et al., 2022). These siderophores can enhance plant Fe acquisition, particularly in Fe-deficient soils, by facilitating the transfer of Fe from the soil to plant roots (Kabir and Bennetzen, 2024; Radzki et al., 2013). Moreover, ethylene-induced flavonoid production is often linked to its role in enhancing root exudation and Fe solubilization under deficiency conditions (Zamioudis et al., 2013). However, the root flavonoid and rhizosphere siderophore levels showed no significant increase upon ACC application in non-stressed soybean plants. The lack of significant increase in root flavonoid and rhizosphere siderophore levels upon ACC application in non-stressed plants suggests that these metabolites are tightly regulated and context-dependent due to the mutual relationship between plant host and microbes. Flavonoid and siderophore production are closely associated with abiotic stress and are predominantly triggered by stress cues rather than ethylene signaling alone (Cesco et al. 2010; Zamioudis and Pieterse, 2012). While ACC supplementation may enhance stress resilience in soybean, its application in control conditions requires caution, as excessive ethylene can act as a growth inhibitor rather than a promoter.

**Figure 8.**
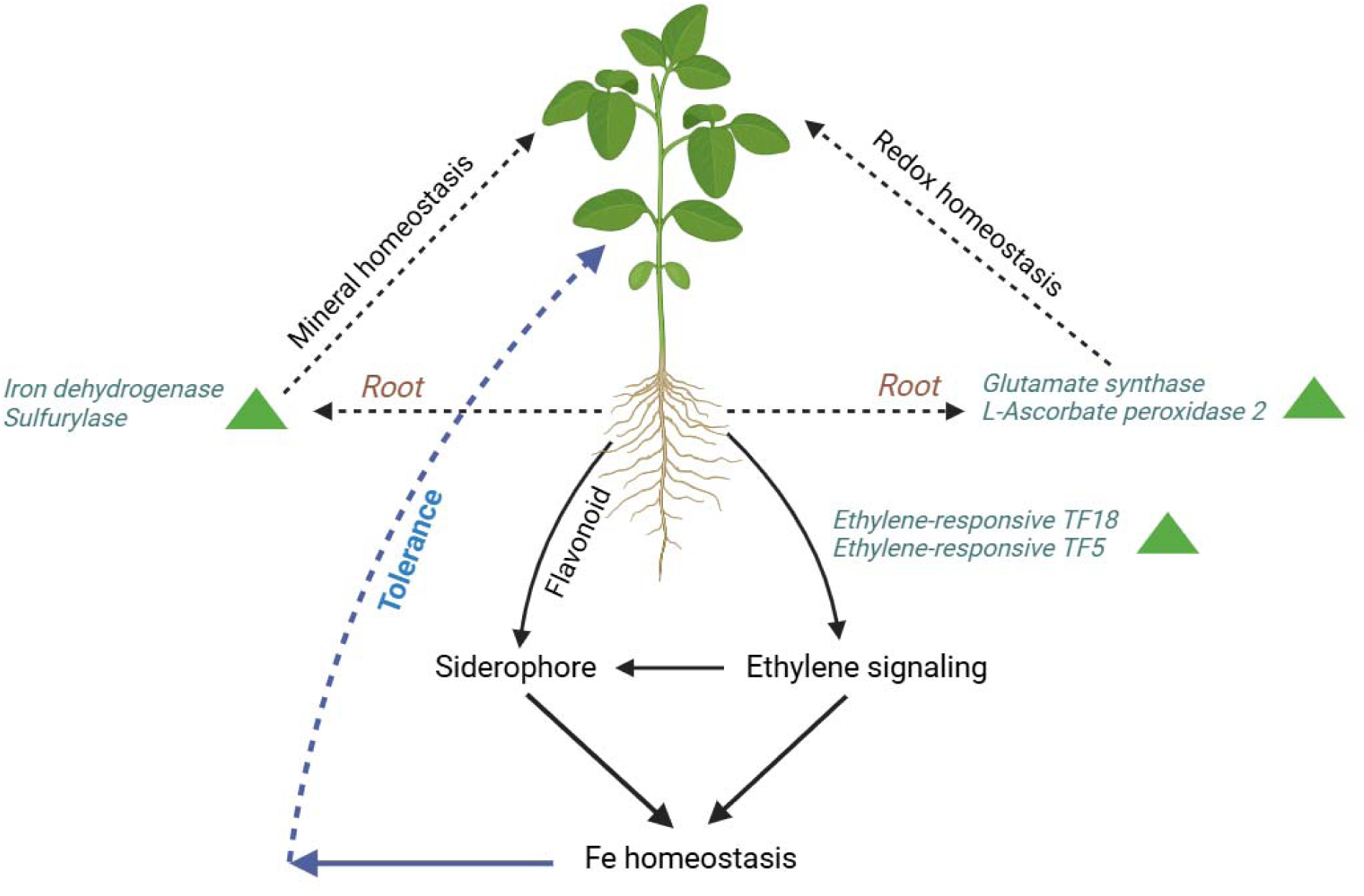
Schematic representation of the molecular and physiological responses of soybean roots under combined soil alkalinity and drought stress. Key processes include mineral homeostasis, redox homeostasis, and ethylene signaling, which collectively contribute to stress tolerance. Genes involved in these pathways, such as *Iron dehydrogenase* and *Sulfurylase* (mineral homeostasis), *Glutamate synthase* and *L-Ascorbate peroxidase 2* (redox homeostasis), and ethylene-responsive transcription factors *TF18* and *TF5* (ethylene signaling), are upregulated in roots. Ethylene signaling modulates root exudation of flavonoids and siderophores, facilitating microbial recruitment to support Fe homeostasis and improve stress resilience. These interactions highlight the coordinated mechanisms promoting soybean adaptation to dual stress conditions.

In this study, the decline in microbial cells due to alkaline and drought stress could be attributed to the combined stress effects on soil and plant environments. Soil alkalinity reduces the availability of essential nutrients and impacts plant root exudation, which serves as a primary energy source for microbes (Chen and Liu, 2024; Pascale et al., 2020). Additionally, drought conditions limit soil moisture, further restricting microbial activity and growth due to reduced mobility and access to nutrients. Furthermore, drought conditions may limit soil moisture, further restricting microbial activity and growth due to reduced mobility and access to nutrients (Qu et al., 2024; Bhattacharyya et al. 2024). Interestingly, we observed the restoration of the overall 16S bacterial and ITS fungal abundance when ACC was augmented in soybeans exposed to dual stresses, compared to plants exposed to Fe-drought+ conditions. Thus, it appears that ethylene signaling may indirectly promote microbial symbiosis, thereby recovering physiological status and drought tolerance in soybean exposed to combined soil alkalinity and drought. Previous studies demonstrated that ethylene plays a complex yet crucial role in enhancing microbial symbiosis in plants by modulating defense responses and facilitating beneficial interactions (Shekhawat et al., 2022; Ravanbakhsh et al., 2018). In legume-rhizobia symbiosis, ethylene levels influence root nodule formation essential for nitrogen fixation (Guinel, 2015). Similarly, in mycorrhizal associations, ethylene can modulate root architecture and colonization (Khatabi et al., 2012). Also, plants exposed to abiotic stresses often produce elevated levels of ethylene to influence root architecture and the composition of root exudates (Glick, 2014). These exudates, which include organic acids, amino acids, and secondary metabolites, act as signaling molecules and attractants for beneficial microbes, such as rhizobacteria and mycorrhizal fungi (Vacheron et al., 2013). Thus, our findings imply that the induction of ethylene-responsive genes may act as a central regulator, integrating signals from both alkaline and drought stress to orchestrate an adaptive response in soybean. This study may encourage future research to uncover the crosstalk between ethylene signaling and microbiome underlying stress resilience in soybean.

The combined soil alkalinity and drought caused downregulation of several genes associated mineral and solute transport, redox homeostasis, and pathogen immunity. Soil alkalinity and/or drought can cause significant nutritional imbalances in plants, disrupting essential metabolic processes and leading to stunted growth and poor health (Kumari et al., 2022). In this study, the downregulation of several genes such as *Anion transporter 6*, *Sulfate transporter 3.4* and *Copper transporter* suggests a possible disruption in nutrient uptake and translocation in soybean exposed to soil alkalinity and drought. Our gene enrichment analysis also indicates the downregulation of ferric Fe binding, Fe-S cluster binding, and metal ion binding, highlighting the critical role of nutrient status in enabling soybean plants to cope with the combined stresses of alkalinity and drought. Drought can cause a decrease in the activity and functionality of anion transporters in plant plasma membranes (Singh et al. 2021b; Jarzyniak and Jasiński, 2014). Soybean plants exhibited downregulation of genes associated with solute and amino acid transport, as observed with *Glyma.10G146600* (solute carrier family 40) and *Glyma.06G222400* (amino acid transporter) due to dual stress. Solute transporters play a critical role in nutrient uptake and ion balance, ensuring the availability of essential minerals for metabolic processes (Sun et al. 2024). Similarly, amino acid transporters are crucial for stress signaling and osmotic adjustment in plants (Delrot et al. 2000). Hence, the lower expression of these transporters limits the ability of soybean to redistribute amino acids for protein metabolism and stress responses, thereby weakening stress adaptation mechanisms. The growth retardation in soybean was further correlated with the downregulation of several genes involved in redox homeostasis which is consistent with the increased ROS exposed to concurrent soil alkalinity and drought. Interestingly, we found that the *Thaumatin* and *Thioredoxin H1* genes, known for their anti-pathogen activity, were suppressed under combined alkalinity and drought stress in soil. Both *Thaumatin* and *Thioredoxin H1* are well-documented for their roles in plant immunity, particularly in mitigating pathogen attacks through anti-fungal and redox-regulating activities (Jedelská et al. 2020; Mata-Pérez et al. 2019). This may be correlated with the reduced microbial dynamics due to soil alkalinity and drought, possibly consisting of beneficial microbes that possess anti-pathogenic activity. In this study, the downregulation of crucial genes reflects a trade-off where plants conserve energy by suppressing nutrient transport and focusing resources on survival, but it also highlights the need for strategies to enhance the expression of these transporters to improve stress resilience in soybean.

## 5 Conclusion

This study emphasizes the impact of soil alkalinity and drought stress on soybean health, with significant reductions in growth and nutrient uptake, although the photochemical efficiency of photosystem II remained unaffected. RNA-seq analysis revealed significant shifts in gene expression, including the upregulation of genes associated with mineral homeostasis, auxin signaling, and reactive oxygen species scavenging, as well as ethylene-responsive transcription factors. Supplementation with an ethylene precursor improved growth, photosynthesis, and Fe status under stress conditions accompanied by elevated root flavonoid and rhizosphere siderophore (Figure 8). However, in non-stressed plants, it caused growth inhibition and reduced Fe levels, underscoring the context-dependent role of ethylene. Furthermore, the ethylene precursor increased root flavonoid and siderophore production while restoring microbial populations under stress, but these effects were reversed in healthy plants. These findings emphasize the critical role of ethylene in regulating root exudates and microbial recruitment to mitigate stress. This study provides valuable insights into leveraging candidate genes and ethylene signaling as strategies to enhance soybean resilience against combined soil alkalinity and drought stress.

## Data availability statement

Raw RNA-seq data was submitted to NCBI under the following accession number: PRJNA1140285.

## Author contributions

MRH cultivated the plants, conducted most of the experiments, performed RNA-seq analysis, and prepared the draft manuscript. AT extracted RNA from root samples and assisted with plant phenotyping. MGM helped with data interpretations and revised the manuscript. AHK conceived the idea, supervised the whole work, and critically revised the manuscript.

## Acknowledgments

This research was supported by a startup grant (5SFAES-293007) and Elizabeth and Hayden Cutler’s Endowed Professorship in Biotechnology from the University of Louisiana at Monroe.

